# Nanopore sequencing for the detection and the identification of *Xylella fastidiosa* subspecies and sequence types from naturally infected plant material

**DOI:** 10.1101/810648

**Authors:** Luigi Faino, Valeria Scala, Alessio Albanese, Vanessa Modesti, Alessandro Grottoli, Nicoletta Pucci, Alessia L’Aurora, Massimo Reverberi, Stefania Loreti

## Abstract

*Xylella fastidiosa* (*Xf*) is a polyphagous gram-negative bacterial plant pathogen that can infect more than 300 plant species. It is endemic in America while, in 2013, *Xf* subsp. *pauca* was for the first time reported in Europe on olive tree in the Southern Italy. The availability of fast and reliable diagnostic tools is indispensable for managing current and future outbreaks of *Xf.*

In this work, we used the Oxford Nanopore Technologies (ONT) device MinION platform for detecting and identifying *Xf* at species, subspecies and Sequence Type (ST) level straight from infected plant material. The study showed the possibility to detect *Xf* by direct DNA sequencing and identify the subspecies in highly infected samples. In order to improve sensitivity, Nanopore amplicon sequencing was assessed. Using primers within the set of the seven MLST officially adopted for identifying *Xf* at type strain level, we developed a workflow consisting in a multiple PCR and an *ad hoc* pipeline to generate MLST consensus after Nanopore-sequencing of the amplicons. The here-developed combined approach achieved a sensitivity higher than real-time PCR allowing within few hours, the detection and identification of *Xf* at ST level in infected plant material, also at low level of contamination.

**Originality Significance Statement:** In this work we developed a methodology that allows the detection and identification of *Xylella fastidiosa* in plant using the Nanopore technology portable device MinION. The approach that we develop resulted more sensitive than methods currently used for detecting *X. fastidiosa*, like real-time PCR. This approach can be extensively used for *X. fastidiosa* detection and it may pave the road for the detection of other tedious vascular pathogens.

## Introduction

*Xylella fastidiosa (Xf)*, a gram-negative phytopathogenic bacterium (Wells *et al.*, 1987), has a very broad host range, causing different diseases in important crops (Hopkins, 1989) and in many urban shade trees (Sherald and Kostka, 1992). *Xf* symptoms may not be evident and many hosts are symptomless making this pathogen very difficult to manage. The pathogen is transmitted by xylem sap feeding insects and colonizes the host xylem vessel, causing the typical leaf scorching. *Xf* is an endemic pathogen in America and only recently was identified in southern Italy on olive trees (Saponari *et al.*, 2013), in Corsica and in the south-east Mediterranean coast of France on several hosts (Denancé *et al.*, 2019). Subsequently, records of *Xf* were reported in Germany on oleander (EPPO, 2016a) and in Spain, Mallorca Island and Andalusia, on different plant species (Landa, 2017; Olmo *et al.*, 2017). Recently, a new finding was also reported in Tuscany (Marchi *et al.*, 2018). Different *Xf* subspecies were identified in Europe in association to the above-mentioned natural outbreaks and interceptions on ornamental plants. There are three formally accepted subspecies of *Xf*, i.e. *fastidiosa, pauca* and *multiplex*, based on DNA–DNA hybridization, as recently confirmed by comparative genomic analysis (Schaad *et al.*, 2002; Marcelletti and Scortichini, 2016; Denancé *et al.*, 2019). However, the International Society of Plant Pathology Committee on the Taxonomy of Plant Pathogenic Bacteria (ISPP-CTPPB) considered valid names only the subsp. *fastidiosa* and subsp. *multiplex* whereas other authors classified five subspecies (*fastidiosa, pauca, multiplex, sandyi* and *morus*) (Bull *et al.*, 2012; Nunney *et al.*, 2012). Complicating the whole taxonomic scenario, *Xf* subspecies includes different sequence types (ST) which are determined by Multi Locus Sequence Typing (MLST) analysis based on the Sanger sequencing technology of seven housekeeping gene (Yuan *et al.*, 2010; Nunney *et al.*, 2012; EPPO, 2016b). MLST analysis is recommended for new findings for the undoubtful identification of the subspecies and ST in case of new outbreak or new plant hosts (EPPO, 2016b). Since the isolation of *Xf* is tedious, the MLST analysis can be performed directly from DNA of the host plant; however, low concentration of the bacteria and contaminants derived from the plant material makes the MLST amplification, thus ST determination, extremely challenging.

Currently, next generation sequencing (NGS) platforms represent high throughput technologies which allow obtaining large datasets of genetic information. In biomedical field, sequencing technologies are rapidly being adopted for bacterial outbreak investigations (Faria *et al.*, 2017) and for rapid clinical diagnostics (Votintseva *et al.*, 2017). A combination of whole transcriptome amplification and Nanopore sequencing device has been used to detect ‘*Candidatus* Liberibacter asiaticus’ or plum pox virus in plants and insect vectors (Bronzato Badial *et al.*, 2018). The reliability of Nanopore for the diagnosis of several plant pathogen was demonstrated (Chalupowicz *et al.*, 2019) and a rapid MLST determination method was described for ST estimation of *Klebsiella pneumoniae* isolates by the ONT MinION device (Page and Keane, 2018).

Direct Nanopore sequencing and Nanopore amplicon sequencing of two or seven housekeeping genes were assessed on naturally infected and spiked samples. Additionally, a consensus was generated from all seven MLST using amplicons sequenced by MinION device. Nanopore amplicon sequencing provides the possibility, within few hours, to detect and identify *Xf*, its subspecie and ST with an equal or slightly better sensitivity than the usual methods of detection and identification or *Xylella* spp. such as real-time PCR.

## Results

### Detection and identification of *Xylella fastidiosa* in infected plant material by direct Nanopore sequencing

Direct Nanopore sequencing was assessed to detect *Xf* in naturally infected plants by using the ONT MinION device. The assay was firstly performed on DNA of 21 olive samples (dataset 1) and 7 samples of different plant species infected by *Xf* (dataset 2) for a total of 28 samples. All samples were analysed by real-time PCR which revealed a variable level of infection among samples, e.g. in samples of dataset 1, Ct ranged from 21.28 to 36.17, indicating a difference of about 10^5^ times between the most infected (Olive-1) and the less infected (Olive-11) sample (Tab. 1). The direct Nanopore sequencing was developed in three experiments, the first one included samples Olive-1 to Olive-12 (dataset 1), the second one, samples from Olive-13 to Olive-21 (dataset 1) and the third experiment was composed of samples from different plant species (dataset 2) (Tab. 2). The first flowcell generated ∼3.2 Gb of data for 12 samples in 14 hours of run, the second flowcell produced ∼2.7 Gb for the remaining nine olive samples in ∼30 hours while the last flowcell generated ∼5.5 Gb in ∼30 hours. All these flowcells were stopped before reaching the 48 hours run due to pore saturation. The three subsets of samples were then subjected to independent de-barcoding and analysis (see methods). After demultiplex of the samples, we were able to retain ∼1.3 Gb (∼40% of the total reads) of data from the first subset of 12 samples, ∼800 Mb (∼30% of the total reads) for the second subset while we got ∼2 Gb of data for the samples from other plant species. The amount of data for individual samples ranged from ∼600 Mb in the *Cystus* sp. sample to ∼10 Mb in Olive-17 displaying a high variability between samples although we tried to use the same amount of input gDNA (Tab. 1, 2). Each sample was analysed using a custom python script (called detection_script) (see material and methods) in order to identify *Xf*. The analysis showed that each sample had a similar bacterial composition and that *Xylella* was the most abundant genus (Tab. 1). Twenty-two out of 28 samples showed reads that map uniquely to the genome of *Xylella* spp. It is worth of mention, that all samples with more reads mapping to the *Xf* genome had a low Ct value (< 31), suggesting that *Xf* can be detected only in heavily infected samples with the flowcell throughput achieved in our work (Tab. 1). To better assess the performance of the direct Nanopore DNA sequencing, we prepared 12 samples of uninfected olive DNA amended with a range of known concentrations of DNA of *X. fastidiosa* subspecies *pauca* strain De Donno (CFBP 8402) (*Xfp*). The amount of DNA was next or below the real-time PCR limit of detection, estimated around 10 fg/PCR reaction (Modesti *et al.*, 2017). Three samples (from LOD-10 to LOD-12) were not detected by real-time PCR giving inconsistent results (Tab. 3). A new flowcell was used to sequence these 12 samples generating ∼667 Mb of data in 22 hours. The quantity of data/sample ranged from 45 Mb to 7.7 Mb and *Xfp* was identified in four samples using the detection_script (Tab. 3). These data confirm that Nanopore direct DNA sequencing can reliably detect *Xf* in samples with a high bacterial concentration.

**Table 1).** Direct Nanopore sequencing of DNA of *Xylella fastidiosa* naturally infected olives samples collected in Apulia region (Dataset 1).

| Sample<br>s | Ct<sup>a</sup> | Mbases | #<br>reads | # reads<br>mapped (%)<sup>b</sup> | # reads mapped<br>on Bacteria (%)<sup>c</sup> | # reads mapping to<br>*Xylella fastidiosa* (%)<sup>d</sup> |
| --- | --- | --- | --- | --- | --- | --- |
| Olive-1 | 21.28 | 140.47 | 46,444 | 41,478 (89.3) | 755 (1.8) | 111 (14.7) |
| Olive-2 | 21.44 | 227.67 | 80,039 | 70,081 (87.6) | 1,236 (1.8) | 188 (15.2) |
| Olive-3 | 21.92 | 113.07 | 34,789 | 31,012 (89.1) | 556 (1.8) | 49 (8.8) |
| Olive-4 | 25.7 | 100.87 | 44,308 | 39,342 (88.8) | 539 (1.4) | 5 (0.9) |
| Olive-5 | 25.5 | 62.23 | 33,632 | 28,633 (85.1) | 367 (1.3) | 3 (0.8) |
| Olive-6 | 25.3 | 68.11 | 33,152 | 28,773 (86.8) | 384 (1.3) | 6 (1.6) |
| Olive-7 | 27.85 | 107.05 | 39,608 | 34,853 (88.0) | 537 (1.5) | 9 (1.7) |
| Olive-8 | 27.24 | 95.05 | 33,240 | 29,871 (89.9) | 525 (1.8) | 2 (0.4) |
| Olive-9 | 27.74 | 102.02 | 39,916 | 35,393 (88.7) | 580 (1.6) | 12 (2.1) |
| Olive-10 | 35.3 | 43.94 | 17,468 | 15,383 (88.1) | 213 (1.4) | 0 (0.0) |
| Olive-11 | 36.17 | 160.30 | 52,157 | 46,588 (89.3) | 827 (1.8) | 0 (0.0) |
| Olive-12 | 35.06 | 43.47 | 18,259 | 15,689 (85.9) | 249 (1.6) | 0 (0.0) |
| Olive-13 | 28 | 69.15 | 30,135 | 25,379 (84.2) | 325 (1.3) | 4 (1.2) |
| Olive-14 | 26 | 111.75 | 42,884 | 37,284 (86.9) | 538 (1.4) | 61 (11.3) |
| Olive-15 | 34.5 | 87.21 | 34,260 | 29,717 (86.7) | 448 (1.5) | 1 (0.2) |
| Olive-16 | 33.75 | 94.79 | 42,478 | 36,246 (85.3) | 480 (1.3) | 0 (0.0) |
| Olive-17 | 31.6 | 31.19 | 10,868 | 9,586 (88.2) | 131 (1.4) | 0 (0.0) |
| Olive-18 | 30.1 | 101.70 | 43,913 | 37,632 (85.7) | 475 (1.3) | 3 (0.6) |
| Olive-19 | 25.3 | 81.98 | 28,591 | 24,781 (86.7) | 402 (1.6) | 22 (5.5) |
| Olive-20 | 36 | 128.11 | 37,611 | 33,844 (90.0) | 543 (1.6) | 0 (0.0) |
| Olive-21 | 29 | 106.21 | 35,496 | 31,236 (88.0) | 540 (1.7) | 4 (0.7) |

<sup>a</sup> Cycle threshold obtained by Harper *et al.*, 2010, TaqMan version
<sup>b</sup> Percentage of reads mapped to bacterial genomes and *Oleae europaea* over the number of total reads sequenced
<sup>c</sup> Percentage of reads mapped to bacterial genomes and *Oleae europaea* from the total mapped
<sup>d</sup> Percentage of reads mapped on *Xylella fastidiosa* genomes over the number of reads mapped to Bacterial genomes

**Table 2).** Direct Nanopore sequencing of DNA of *Xylella fastidiosa* naturally infected plant samples collected from Tuscany Region (dataset 2)

| Samples | Ct<sup>a</sup> | Mbases | # reads | # reads mapped (%)<sup>b</sup> | # reads mapped on Bacteria (%)<sup>c</sup> | # reads mapping to *Xylella fastidiosa* (%)<sup>d</sup> |
| --- | --- | --- | --- | --- | --- | --- |
| *Prunus dulcis* | 27,35 | 122.78 | 114,327 | 11,508 (10.1) | 7,621 (66.2) | 22 (0.3) |
| *Rhamnus alaternus* | NA | 73.45 | 62,664 | 4,335 (6.9) | 2,737 (63.1) | 10 (0.4) |
| *Cystus* | 25,78 | 606.32 | 199,423 | 5,156 (2.6) | 3,522 (68.3) | 230 (6.5) |
| *Rosmarinus officinalis* | 35,14 | 247.79 | 182,831 | 9,657 (5.3) | 4,662 (48.3) | 2 (0.1) |
| *Lavandula* | 28,75 | 438.39 | 211,320 | 25,197 (11.9) | 10,231 (40.6) | 86 (0.8) |
| *Polygala myrtifolia* | 25,10 | 184.91 | 87,762 | 11,447 (13.0) | 6,157 (53.8) | 104 (1.7) |
| *Spartium juceum* | 18,61 | 286.19 | 155,150 | 10,919 (7.0) | 6,650 (60.9) | 1,699 (25.5) |

<sup>a</sup> Cycle threshold obtained by Francis *et al.*, 2006, TaqMan version
<sup>b</sup> Percentage of reads mapped to bacterial genomes and *Oleae europaea* over the number of total reads sequenced
<sup>c</sup> Percentage of reads mapped to bacterial genomes and *Oleae europaea* from the total mapped
<sup>d</sup> Percentage of reads mapped on *Xylella fastidiosa* genomes over the number of reads mapped to bacterial genomes

### Nanopore amplicon sequencing

The MinION device generated about 290 Mb of data for all 12 samples (dataset 3) and after reads demultiplex, we retained ∼155 Mb of data. The number of reads is in a range of 14,360 and 54,405 for LOD-10 and LOD-2, respectively (Tab. 4). The detection_script was used to detect *Xf* in all samples. The results showed that the number of reads matching uniquely a bacterial genome varied between 3.39% in sample LOD-9 and 50.53% in sample LOD-5 (Tab. 4). As expected, the most abundant genera represented in the mapped reads, >95% of the reads mapped to *Xf* (Tab. 4, Fig. 1). The only exception is the sample LOD-9 for which low number of reads mapped to bacterial genomes suggesting a problem in the sample preparation (Tab. 4, Fig. 1). The reduction of sequencing error – since the same region is sequenced more than once - produces a more accurate output allowing us to identify *Xf* at the subspecies level. To validate this, we used the detection_script in combination with a specific *Xf* database formed by every *Xf* genome available at the NCBI database (∼60 genomes). The results showed that all 11 samples positive for *Xf* had >95% of the reads correctly mapped to a genome identified as *Xf* subsp. *pauca* which is the subspecies amended to the olive gDNA (Fig. 2). The same approach was used – amplification of *cysG* and *malF*, Nanopore sequencing, querying the *Xf*-customised database – with the dataset 2 composed of gDNA from different *Xf* naturally infected plant species (Tab. 5, Fig. 3). The results showed that for all samples, except for *Rhamnus alaternus*, more than half of the reads mapped to *Xf*. Additionally, the analysis using the specific *Xf* database showed that >90% of the reads mapped to *Xf* subsp. *multiplex* genome (Fig. 3). To further support the subspecies identification results in the unknown samples, we generated a consensus for the *cysG* and *malF* sequences using the consensus_script. These consensus sequences were aligned to the *Xf* specific database matching 100% to *multiplex* subspecies. This device correctly identify the *Xf* subspecie *multiplex* associated to the Tuscany samples, according to the recent report of Marchi *et al.* (2018).

**Figure 1).**
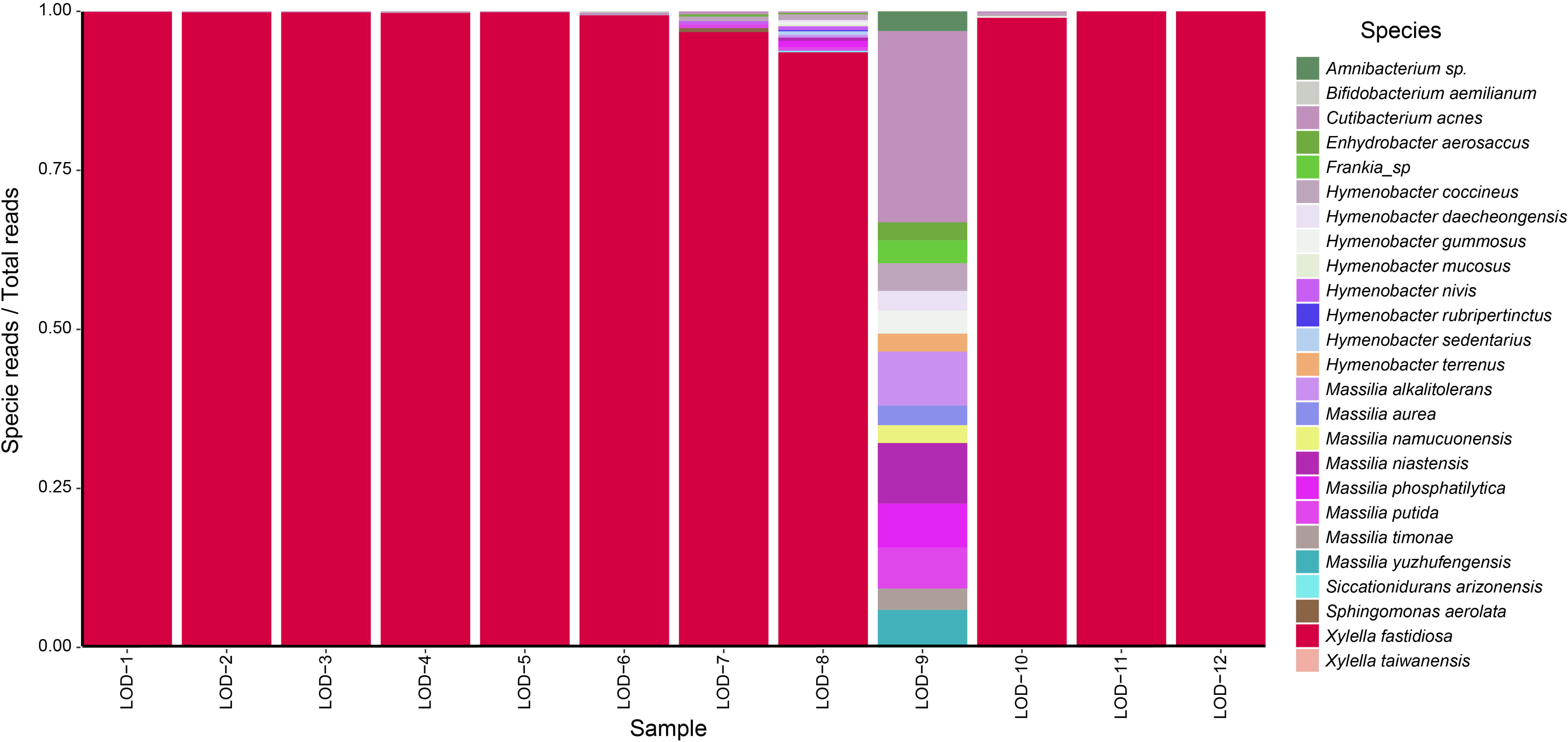
Stacked bar charts showing the actual relative abundance of bacterial families in the identified using the detection_script on Nanopore ampliseq of olive DNA amended with different amount of *X. fastidiosa* subspecie *pauca* DNA

**Figure 2).**
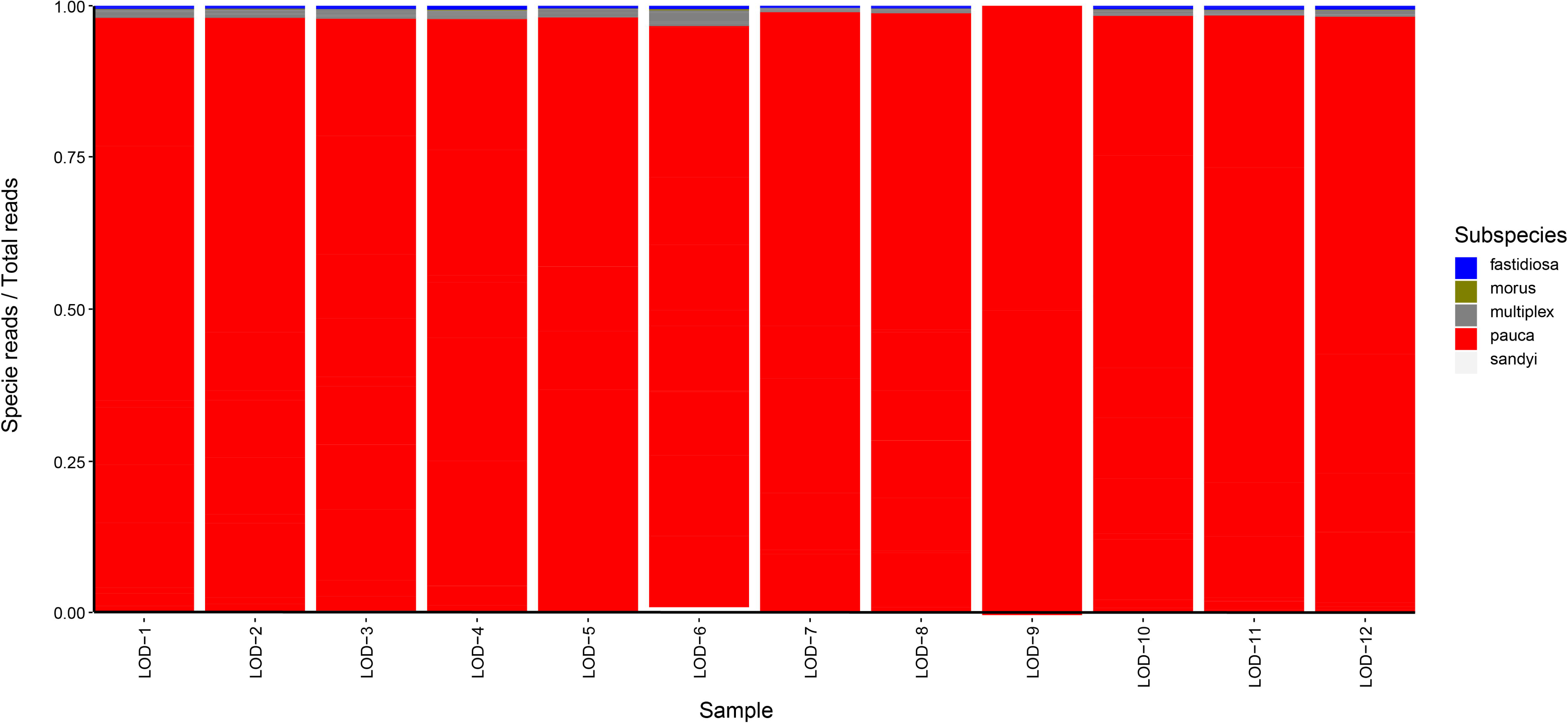
Stacked bar charts showing the actual relative abundance of *X. fastidiosa* subspecie *pauca* compared to other *X. fastidiosa* subspecies using the detection_script on Nanopore ampliseq of olive DNA amended with different amount of *X. fastidiosa* subspecie *pauca* DNA

**Figure 3).**
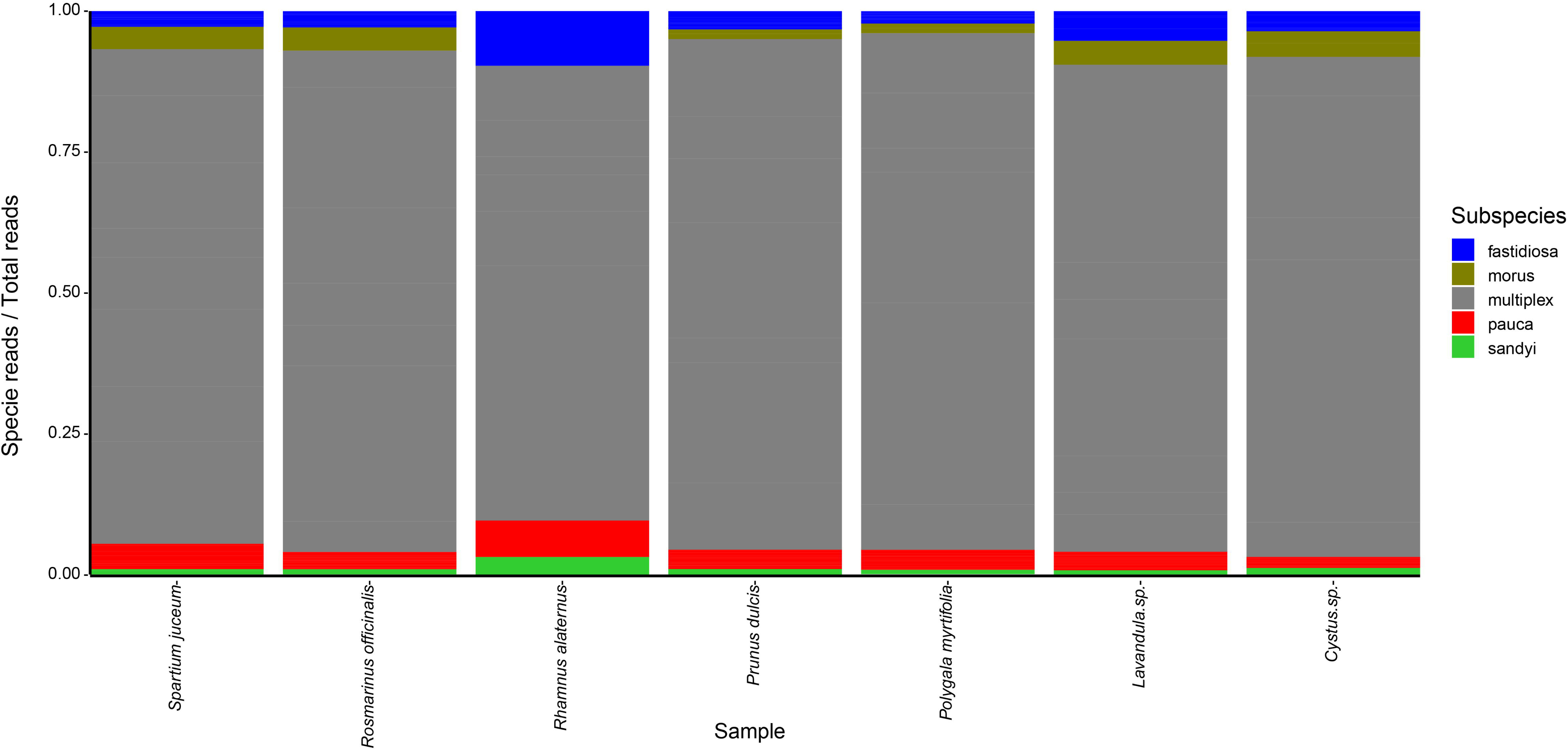
Stacked bar charts showing the actual relative abundance of different *X. fastidiosa* subspecie using the detection_script on Nanopore ampliseq of DNA from different plant species. The highest subspecies present resulted to the *X. fastidiosa* subspecies *multiplex*

**Table 4).**
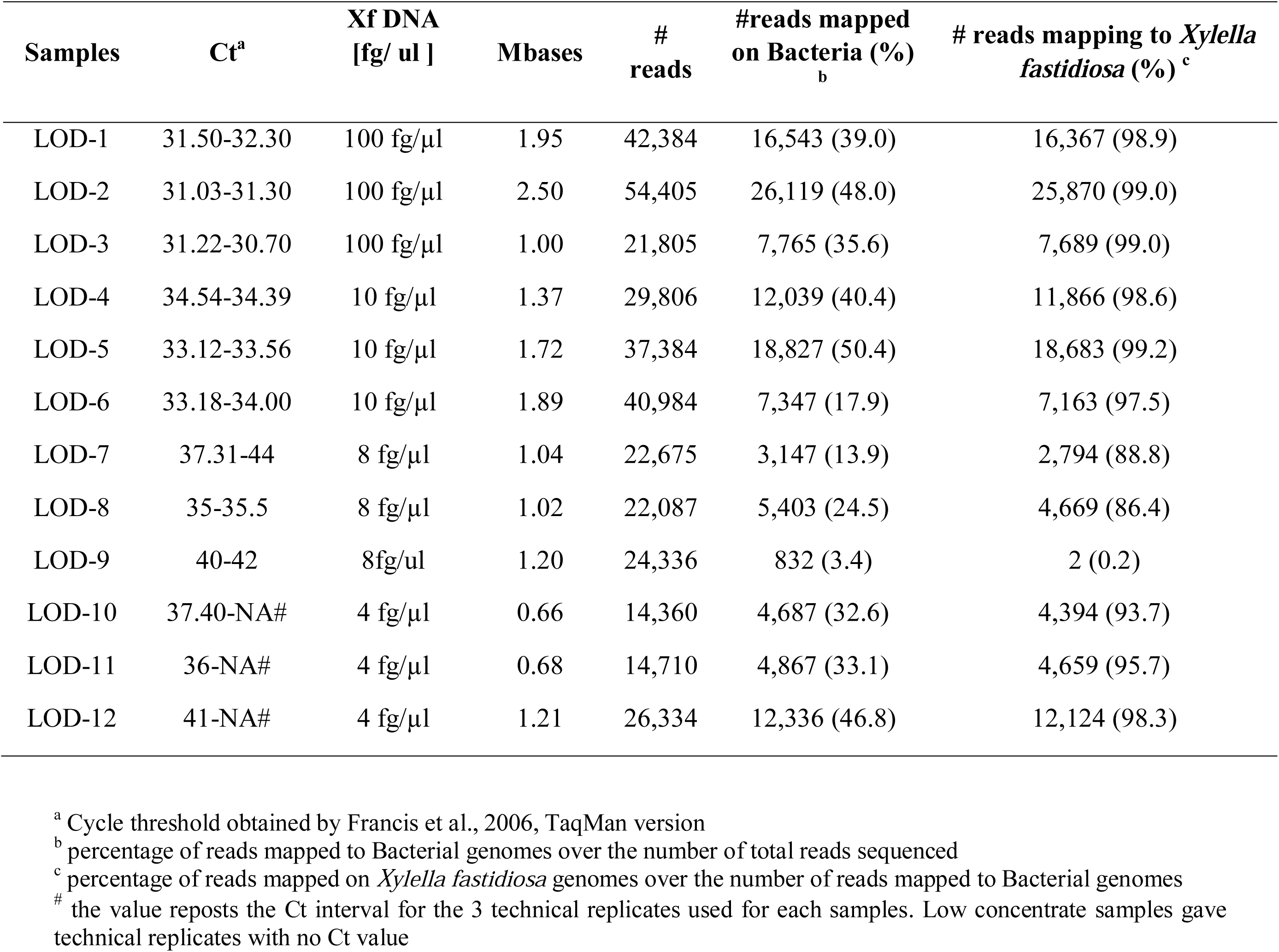
Nanopore amplicon sequencing of *cysG* and *malF* from DNA of healthy olives samples spiked with known amount of *Xylella fastidiosa* subsp. *pauca* strain De Donno (CFBP 8402) DNA (Dataset 3)

| Samples | Ct<sup>a</sup> | Xf DNA<br>[fg/ ul ] | Mbases | #<br>reads | #reads mapped<br>on Bacteria (%)<br><sub>b</sub> | # reads mapping to *Xylella*<br>*fastidiosa* (%) <sup>c</sup> |
| --- | --- | --- | --- | --- | --- | --- |
| LOD-1 | 31.50-32.30 | 100 fg/μl | 1.95 | 42,384 | 16,543 (39.0) | 16,367 (98.9) |
| LOD-2 | 31.03-31.30 | 100 fg/μl | 2.50 | 54,405 | 26,119 (48.0) | 25,870 (99.0) |
| LOD-3 | 31.22-30.70 | 100 fg/μl | 1.00 | 21,805 | 7,765 (35.6) | 7,689 (99.0) |
| LOD-4 | 34.54-34.39 | 10 fg/μl | 1.37 | 29,806 | 12,039 (40.4) | 11,866 (98.6) |
| LOD-5 | 33.12-33.56 | 10 fg/μl | 1.72 | 37,384 | 18,827 (50.4) | 18,683 (99.2) |
| LOD-6 | 33.18-34.00 | 10 fg/μl | 1.89 | 40,984 | 7,347 (17.9) | 7,163 (97.5) |
| LOD-7 | 37.31-44 | 8 fg/μl | 1.04 | 22,675 | 3,147 (13.9) | 2,794 (88.8) |
| LOD-8 | 35-35.5 | 8 fg/μl | 1.02 | 22,087 | 5,403 (24.5) | 4,669 (86.4) |
| LOD-9 | 40-42 | 8fg/ul | 1.20 | 24,336 | 832 (3.4) | 2 (0.2) |
| LOD-10 | 37.40-NA# | 4 fg/μl | 0.66 | 14,360 | 4,687 (32.6) | 4,394 (93.7) |
| LOD-11 | 36-NA# | 4 fg/μl | 0.68 | 14,710 | 4,867 (33.1) | 4,659 (95.7) |
| LOD-12 | 41-NA# | 4 fg/μl | 1.21 | 26,334 | 12,336 (46.8) | 12,124 (98.3) |

<sup>a</sup> Cycle threshold obtained by Francis et al., 2006, TaqMan version
<sup>b</sup> percentage of reads mapped to Bacterial genomes over the number of total reads sequenced
<sup>c</sup> percentage of reads mapped on *Xylella fastidiosa* genomes over the number of reads mapped to Bacterial genomes
<sup>#</sup> the value reposts the Ct interval for the 3 technical replicates used for each samples. Low concentrate samples gave technical replicates with no Ct value

**Table 5).** Nanopore-amplicon-sequencing of *cysG* and *malF* from DNA of naturally infected plants samples collected in Tuscany region

| Samples | Ct<sup>a</sup> | Mbases | # reads | # reads mapping to *Xylella fastidiosa* (%)<sup>b</sup> |
| --- | --- | --- | --- | --- |
| *Prunus dulcis* | 27,35 | 0.28 | 6,108 | 3,396 (55.6) |
| *Rhamnus alaternus* | NA | 0.44 | 9,462 | 28 (0.3) |
| *Cistus* | 25,78 | 0.44 | 9,600 | 5,656 (58.9) |
| *Rosmarinus officinalis* | 35,14 | 0.29 | 6,228 | 2,178 (35.0) |
| *Lavandula* | 28,75 | 1.19 | 25,798 | 10,391 (40.3) |
| *Polygala myrtifolia* | 25,10 | 0.51 | 11,085 | 7,065 (63.7) |
| *Spartium jaceum* | 18,61 | 0.50 | 10,959 | 5,633 (51.4) |

<sup>a</sup> Cycle threshold obtained by Francis et al., 2006, TaqMan version
<sup>b</sup> percentage of reads mapped on *Xylella fastidiosa* genomes over the number of total reads sequenced

### Identification of *Xylella* sequence-type using Nanopore amplicon sequencing

MLST consensus for all seven sequences for three (*Cystus* sp., *Rosmarinus officinalis, Lavandula* sp.) out of the seven samples within the dataset 3 was generated using both Sanger and Nanopore sequencing. These plant species were selected because the ST of *Xf* had not already been determined. The consensus_script was used to generate a consensus using only Nanopore errors-prone reads. As expected, the script generated seven sequences for each sample that were compared with the same sequences generated by Sanger technology (Yuan *et al.*, 2010) followed by querying the database at htpp://pubmlst.org/xfastidiosa/ (Jolley *et al.*, 2018). Pairwise alignment of concatenated sequences of all sever MLST derived from Sanger and Nanopore consensus resulted in a 100% identity (Fig. 4). Additionally, when was compared the concatenated sequences from both Sanger and Nanopore consensus sequences to the *Xf* MLST database, was found that samples of *Cystus* sp., *Rosmarinus officinalis* and *Lavandula* sp. were infected by the ST 87 (Fig. 4). This result is in agreement with recent findings of *Xf* ST 87 in *Prunus amygdalus, Polygala myrtifolia, Spartium junceum* and *Rhamnus alaternus* (Saponari *et al.*, 2019).

**Figure 4).**
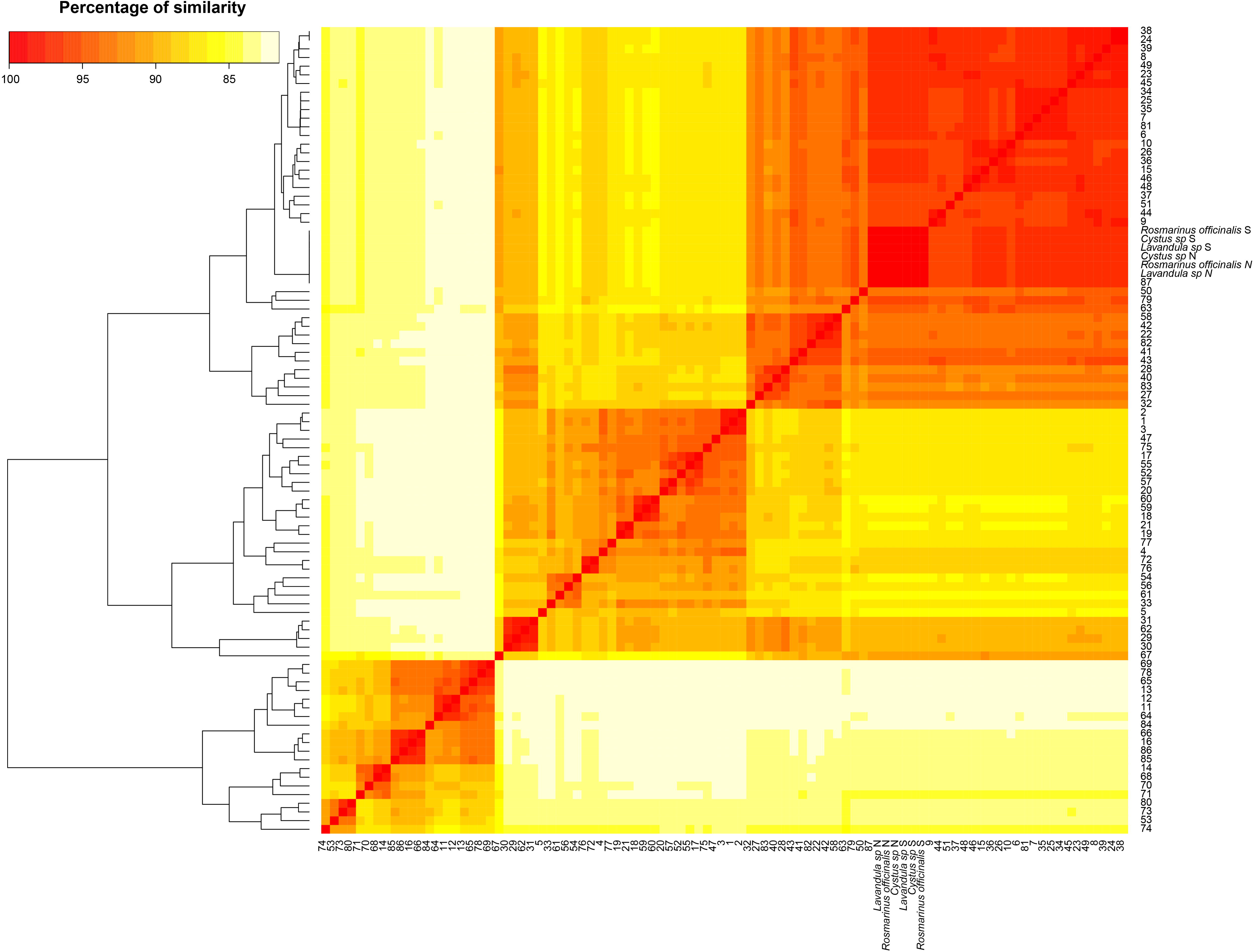
Heatmap representing sequence alignments of seven concatenated MLST for 87 ST deposited at the MLST database (https://pubmlst.org), three sequences derived from Sanger sequencing of *Rosmarinus officinalis, Cystus* sp. and *Lavandula* sp. and three sequences derived from consensus of the Nanopore sequencing for *Rosmarinus officinalis, Cystus* sp. and *Lavandula* sp. plant samples

## Discussion

In the last fifteen years, NGS transformed genomic research with an inevitable impact also on diagnostics, taking us to a new scenario. These technologies have recently been applied for the diagnosis of plant pathogens (Bronzato Badial *et al.*, 2018; Chalupowicz *et al.*, 2019) and for the detection and identification of *Xf* (Bonants *et al.*, 2019). Although several diagnostic methods are available for the detection of *Xf* (EPPO, 2016b), the identification of subspecies and ST are currently laborious and time-consuming. Recently, a tetraplex qPCR assays was developed for simultaneous detection and identification of subspecies in plant tissues (Enora *et al.*, 2019). The detection of *Xf* at subspecies level is fundamental and in case of new outbreaks or new plant hosts, the identification of the ST is strongly recommended (EPPO Standard PM7/24 (4)). To circumvent the complexity of the detection and identification of *Xf* subspecies and ST, we investigate the use of Oxford Nanopore Technologies (ONT) MinION device.

Direct Nanopore DNA sequencing was firstly assessed by using naturally infected samples (dataset 1 and 2). One of the advantages of MinION device compared to other NGS platform is the portability.

However, our results showed that only in highly infected samples *Xf* can be reliably detected and its subspecies identified. All samples lacking reads mapping to the *Xf* genomes, showed a high Ct value by real-time PCR that, associated to low throughput for these samples, made *Xf* undetectable. This evidence was confirmed by using artificially spiked olive samples with DNA concentration of *Xf* next and below the limit of detection of the real-time PCR (dataset 3), currently considered the most sensitive assay for *Xf* detection (EPPO, 2016b; Modesti *et al.*, 2017). Nanopore direct DNA sequencing of these samples provides a very low number of reads which reflect an even lower number of reads mapping to *Xf* genomes making the analysis doubtable. A similar result was also obtained by Bonants *et al.* (2019) using Illumina, who reported the ability of NGS to determine the ST of *Xf* only in highly infected samples. The reason of the low sensitivity of the direct Nanopore sequencing should be addressed to the low quality of the DNA for the tested samples. Nanopore sequencing suffer of low throughput when the DNA used for the sequencing has contaminants and/or when reads are shorts (Chalupowicz *et al.*, 2019). Further optimization of the DNA purification and size selection step may be necessary to maximize the performance of the Nanopore system. This aspect is of relevant importance because *Xf* has a very broad host-range and even using the same extraction method, the DNA quality can be compromised by contaminants present in the host matrix. However, carrying out massive analyzes cannot provide time-consuming or expensive DNA purifications and generally neither the DNA concentration nor the DNA quality are determined before a routinely analysis. Charalampous *et al*. (2019) developed a DNA extraction in order to deplete host DNA from the samples and have more pathogen DNA. Using this approach, they were able to identify respiratory pathogens in human samples. An alternative to sophisticated DNA extraction methodology is amplicon sequencing (Kilianski *et al.*, 2015; Radhakrishnan *et al.*, 2019). This latter approach was assessed in order to overcome the low throughput of the flowcells and to set up a more reliable and sensitive method, based on the amplification of two housekeeping genes followed by Nanopore sequencing. Sequencing of *cysG* and *malF* for subspecies discrimination is suggested by the EPPO Standard PM7/24 (4) and required, since 2018, in France (Enora *et al.*, 2019). Nanopore amplicon sequencing was more efficacious than real-time PCR in *Xf* detection of the spiked samples with low concentration of *Xf* target DNA (dataset 3, Tab. 3). These results suggest that Nanopore amplicon sequencing has better sensitivity than real-time PCR. The addition of the amplification step, even if lengthening the procedure, showed several advantages: *i)* reduces the influence of DNA quality; *ii)* mitigates sequence error by repeatedly sequencing the same region; *iii*) produces usable data faster than genomic DNA sequencing and *iv)* by using lower stringency condition (40 cycles) in the amplification step, it obtains higher number of copies of the target genes unusable for Sanger sequencing but exploitable for Nanopore sequencing (data not shown).

The addition of the amplification allows in a single sequencing step the detection of the pathogen as well as the identification of its subspecies and ST. The results of amplicon sequencing of the dataset 2 (Tab. 2) (*Spartium junceum, Polygala myrtifolia, Rosmarinus offcinalis, Lavandula, Prunus amygdalus, Cystus*) confirmed previous findings which identified as *multiplex* the subspecies infecting these plants (Marchi *et al.*, 2018; Saponari *et al.*, 2019). These evidences showed that by using an amplification step the method was able to detect and identify *Xf* in different plant species, without interference due to the DNA quality.

The results obtained by amplicon sequencing of cysG and malF encourage us to test all the seven housekeeping genes (leuA, petC, malF, cysG, holC, nuoL, gltT) to define the ST in our naturally infected samples. For this purpose, Nanopore and Sanger sequencing of the seven housekeeping genes were performed in Lavandula sp., Rosmarinus officinalis and Cystus sp. collected in Tuscany Region, for which the ST has not yet been reported. Our results showed that Xf recovered from these samples belong to the new ST 87 accordingly with previous findings on Prunus amygdalus, Polygala myrtifolia, Spartium junceum and Rhamnus alaternus (Saponari et al., 2019). The simultaneous detection and identification of Xf, its subspecie and ST, developed in this study, lead the Nanopore amplicon sequencing assay to be a powerful tool for a quick Xf diagnosis. The evidences obtained in this study shows that the sequencing of two or seven housekeeping genes by MinION is a promising alternative to detect and identify Xf from infected plant material, also in low bacterial concentrations. This higher sensitivity is of interest for Xf detection in traded plants and for latently infected material that represent one of the most serious threat for the dissemination. For an “in field” application, further studies are required to make DNA direct Nanopore sequencing reliable in detecting low concentration pathogens, i.e. using single flow-cells for the processing of individual samples. This allows a greater depth in sequencing, with consequent higher possibility of detection and identification of Xf /subspecies/STs. In conclusion, the evidences obtained in this study paving the way for new opportunities of Nanopore sequencing as an effective survellaince tool for Xf early detection.

## Experimental procedures

### Samples and DNA extraction

#### Sample preparation

Three datasets of samples were prepared as following: the first dataset (1) consists of twenty-one DNA samples extracted from naturally infected olive plants (*Olea europea* L.) collected in Apulia region (southern Italy) as described in Scortichini *et al*. (2018). The second dataset (2) consisted of naturally infected samples of *Spartium junceum, Polygala myrtifolia, Rosmarinus officinalis, Rhamnus alaternus, Prunus amygdalus, Cistus* and *Lavandula* spp., collected in Tuscany. The third dataset (3) consists of DNA of healthy olives tree (collected in Latium) spiked with known quantity of DNA of *Xfp* strain CFBP 8402. Plant DNA extraction of dataset 2 and 3 were performed by CTAB-based method as reported in EPPO Standard PM7/ 24 (4).

A pure colture of *Xfp* strain CFBP 8402 was grown for seven days at 28° C in BYCE medium. The culture scraped and resuspended in 100 μl of PBS, was grown in 10 ml of PD2 broth at 28° C, 170 rpm, for 7-10 days. The bacterial DNA was extracted from 700 µl of pure culture using the Gentra Puregene Yeast/Bact Kit (Qiagen, The Netherlands). DNA (about 40 ng/µl) of healthy olive samples was amended with 100, 10, 8, 4 fg/PCR reaction of *Xfp* DNA, each in three independent replicates.

#### Real-time PCR and Multi-Locus Sequence Typing (MLST)

Datasets DNA were quantified by DS-11 FX+ spectrophotometer (DENOVIX) and diluted to a final concentration of 20 ng/μL. *Xf* was detected in dataset 1 DNA by real-time PCR as described by Harper *et al.* (2010) in a final volume of 10 µl and in datasets 2 and 3 by Francis *et al.* (2006). All samples were also tested by Li *et al.* (2013) to confirm the bacterial infection. MLST analysis was performed on *Cystus, Lavandula* and *Rosmarinus* as previously described (Yuan *et al.*, 2010; EPPO, 2016b). Subspecie identification by sequencing *cysG* and *malF* was performed in dataset 2. For Nanopore amplicon sequencing the amplification of housekeeping genes was modified increasing the cycles of PCR condition from 35 to 40.

#### Datasets used for Nanopore sequencing

Direct Nanopore DNA sequencing was performed on the three previously described datasets (Tab. 1, 2, 3). Dataset 3 was used to test the direct DNA Nanopore sequencing in condition that was next the limit of detection (LoD) of the real-time PCR (Tab. 3). DNA samples from dataset 2 and 3 were used for Nanopore amplicon sequencing of the two genes, *cysG* and *malF*. Three out of seven samples of the dataset 2 (i.e. *Cystus, Lavandula* and *Rosmarinus)* were assessed by Nanopore amplicon sequencing of all seven housekeeping genes.

**Table 3).** Direct Nanopore sequencing of healthy olives samples DNA spiked with known DNA concentrations of *Xylella fastidiosa* subsp. *pauca* strain De Donno (CFBP 8402) DNA (Dataset 3)

| Samples | Ct<sup>a</sup> | *Xf* DNA<br>[fg/μl] | fg gDNA<sup>b</sup> | Mbases | # reads | # reads<br>mapped<br>(%)<sup>c</sup> | #reads<br>mapped on<br>Bacteria (%)<sup>d</sup> | # reads<br>mapping to<br>*Xylella*<br>*fastidiosa*<br>(%)<sup>e</sup> |
| --- | --- | --- | --- | --- | --- | --- | --- | --- |
| LOD-1 | 31.50-32.30 | 100 fg/μl | 700fg | 33.63 | 21,218 | 16,334 (77.0) | 615 (3.8) | 2 (0.3) |
| LOD-2 | 31.03-31.30 | 100 fg/μl | 700fg | 7.97 | 6,656 | 3,853 (57.9) | 105 (2.7) | 0 (0.0) |
| LOD-3 | 31.22-30.70 | 100 fg/μl | 700fg | 17.51 | 16,237 | 8,820 (54.3) | 308 (3.5) | 1 (0.3) |
| LOD-4 | 34.54-34.39 | 10 fg/μl | 70fg | 26.35 | 22,475 | 15,047 (66.9) | 542 (3.6) | 0 (0.0) |
| LOD-5 | 33.12-33.56 | 10 fg/μl | 70fg | 26.31 | 21,669 | 14,915 (68.8) | 486 (3.3) | 1 (0.2) |
| LOD-6 | 33.18-34.00 | 10 fg/μl | 70fg | 31.99 | 24,385 | 16,836 (69.0) | 597 (3.5) | 0 (0.0) |
| LOD-7 | 37.31-44 | 8 fg/μl | 56fg | 27.69 | 21,949 | 14,693 (66.9) | 475 (3.2) | 0 (0.0) |
| LOD-8 | 35-35.5 | 8 fg/μl | 56fg | 35.64 | 34,572 | 23,594 (68.2) | 725 (3.1) | 0 (0.0) |
| LOD-9 | 40-42 | 8fg/ul | 56fg | 22.15 | 29,800 | 18,724 (62.8) | 455 (2.4) | 0 (0.0) |
| LOD-10 | 37.40-NA<sup>#</sup> | 4 fg/μl | 28fg | 39.51 | 38,724 | 26,887 (69.4) | 740 (2.8) | 1 (0.1) |
| LOD-11 | 36-NA<sup>#</sup> | 4 fg/μl | 28fg | 45.19 | 45,824 | 30,053 (65.6) | 833 (2.8) | 0 (0.0) |
| LOD-12 | 41-NA<sup>#</sup> | 4 fg/μl | 28gf | 10.07 | 21,082 | 10,627 (50.4) | 272 (2.6) | 0 (0.0) |

<sup>a</sup> Cycle threshold obtained by Francis *et al.*, 2006, TaqMan version The value reports the Ct interval for the 3 technical replicates used for each sample. Low concentrate samples gave technical replicates with no Ct value
<sup>b</sup> Total amount of *X. fastidiosa* subspecies *pauca* gDNA added to *Oleae europaea* gDNA and used for Nanopore sequencing
<sup>c</sup> Percentage of reads mapped to bacterial genomes and *Oleae europaea* over the number of total reads sequenced
<sup>d</sup> Percentage of reads mapped to bacterial genomes and *Oleae europaea* from the total mapped
<sup>e</sup> Percentage of reads mapped on *Xylella fastidiosa* genomes over the number of reads mapped to bacterial genomes

#### Nanopore sequencing

Nanopore sequencing libraries were prepared according to manufacture instruction for the kit SQK-RBK004 for both direct DNA and amplicon sequencing. In brief, about ∼400 ng of DNA was purified using AMPure beads, ligated to the indexing adapter, combined in one sample and subsequently ligated to the RAP adapter prior sequencing. For amplicon sequencing, PCR amplicons of each sample were pulled together and purified using Isolate II PCR and gel Kit (BIOLINE) and about 100 ng of DNA was used in library preparation. DNA samples were run on the flowcell until pore life ended while amplicon sequencing runs were performed for shorted time (∼5hours). After sequencing, Deepbinner (v0.2.0)(Wick *et al.*, 2018) was used to de-multiples the samples by default parameters. Subsequently, basecalling and a new run of demultiplex was performed using Guppy (v2.3.1; default parameters)(Wick *et al.*, 2019). Deepbinner and guppy basecalling were performed on a GPU card Nvidia GTX 1070 8Gb.

#### Pipeline for gDNA Xylella detection: detection_script

Reads generated by the two-step de-barcoding analysis were analysed using a custom pipeline. In brief, reads were mapped to a database by using Minimap2 (v2.17-r941)((Li, 2018) software using --MD and --secondary=no as additionally parameters. The mapping is split in two steps: the first step aligns all the reads against a database that includes one representative genome for all sequenced bacterial species (about 10,500 bacterial genomes) present at the NCBI Genebank database and a second step that align all reads mapped to *Xf* in the first step to a *Xf* specific database. The *Xf* database it is formed by ∼60 genomes. For both steps, alignment output files are parsed to retrieve only the best match for reads mapping on multiple genomes. Finally, the number of reads mapping on the same subspecies/strain are combine and summarized in plot (https://github.com/lfaino/xylella).

#### MLST consensus generation: consensus_script

A second pipeline was written in order to generate MLST consensus after Nanopore sequencing. Briefly, reads from Nanopore sequencing are demultiplexed by using Deepbinner software (v0.2.0)(Wick *et al.*, 2018). Subsequently, porechop software (v0.2.4) is used to remove adapters from the kept reads. Two rounds of reads correction by racon software (v1.3.3) (Vaser *et al.*, 2017) are performed. The corrected reads are passed to jellyfish software (v2.2.8) (Marçais and Kingsford, 2011) and reads of 100 nt are generated. These reads are assembled by SPADES software (v3.12.0) (Vyahhi *et al.*, 2012) and the assembled contigs polished by Nanopolish (v0.11.1) and subsequently by bcftools mpileup software (v1.7.2) (Danecek and McCarthy, 2017). The reconstructed MLST sequences are compared to all other MLST deposited at https://pubmlst.org/ (Jolley *et al.*, 2018). The script used for MLST reconstruction can be found at GitHub (https://github.com/lfaino/xylella).

## Acknowledgements

This work was supported by MIPAAFT, Project Oli.Di.X.I.It (“OLIvicoltura e Difesa da *Xylella fastidiosa* e da Insetti vettori in Italia”), D.M. 23773 del 6/09/2017.

## References

Bonants, P., Griekspoor, Y., Houwers, I., Krijger, M., van der Zouwen, P., van der Lee, T.A.J., and van der Wolf, J. (2019) Development and Evaluation of a Triplex TaqMan Assay and Next-Generation Sequence Analysis for Improved Detection of *Xylella* in Plant Material. Plant Dis 103: 645–655.

Bronzato Badial, A., Sherman, D., Stone, A., Gopakumar, A., Wilson, V., Schneider, W., and King, J. (2018) Nanopore Sequencing as a Surveillance Tool for Plant Pathogens in Plant and Insect Tissues. Plant Dis 102: 1648–1652.

Bull, C.T., De Boer, S.H., Denny, T.P., Firrao, G., Fischer-Le Saux, M., Saddler, G.S., et al. (2012) List of new names of plant pathogenic bacteria (2008-2010). J Plant Pathol 94: 21–27.

Chalupowicz, L., Dombrovsky, A., Gaba, V., Luria, N., Reuven, M., Beerman, A., et al. (2019) Diagnosis of plant diseases using the Nanopore sequencing platform. Plant Pathol 68: 229–238.

Charalampous, T., Kay, G.L., Richardson, H., Aydin, A., Baldan, R., Jeanes, C., et al. (2019) Nanopore metagenomics enables rapid clinical diagnosis of bacterial lower respiratory infection. Nat Biotechnol 37: 783–792.

Danecek, P. and McCarthy, S.A. (2017) BCFtools/csq: haplotype-aware variant consequences. Bioinformatics 33: 2037–2039.

Denancé, N., Briand, M., Gaborieau, R., Gaillard, S., and Jacques, M.A. (2019) Identification of genetic relationships and subspecies signatures in *Xylella fastidiosa*. BMC Genomics 20:.

Enora, D., Martial, B., Marie-Agnès, J., and Sophie, C. (2019) Novel tetraplex qPCR assays for simultaneous detection and identification of *Xylella fastidiosa* subspecies in plant tissues. bioRxiv 699371.

EPPO (2016a) First report of *Xylella fastidiosa* in Spain. EPPO Report Serv 11: 133.

EPPO (2016b) PM 3/81 (1) Inspection of consignments for *Xylella fastidiosa*. EPPO Bull Stand 46: 395–406.

Faria, N.R., Quick, J., Claro, I.M., Theze, J., de Jesus, J.G., Giovanetti, M., et al. (2017) Establishment and cryptic transmission of Zika virus in Brazil and the Americas. Nature 546: 406.

Francis, M., Lin, H., Rosa, J.C.-L., Doddapaneni, H., and Civerolo, E.L. (2006) Genome-based PCR primers for specific and sensitive detection and quantification of *Xylella fastidiosa*. Eur J Plant Pathol 115: 203–213.

Harper, S.J., Ward, L.I., and Clover, G.R.G. (2010) Development of LAMP and Real-Time PCR Methods for the Rapid Detection of *Xylella fastidiosa* for Quarantine and Field Applications. Phytopathology 100: 1282–1288.

Hopkins, D.L. (1989) *Xylella Fastidiosa*: Xylem-Limited Bacterial Pathogen of Plants. Annu Rev Phytopathol 27: 271–290.

Jolley, K.A., Bray, J.E., and Maiden, M.C.J. (2018) Open-access bacterial population genomics: BIGSdb software, the PubMLST. org website and their applications. Wellcome open Res 3:.

Kilianski, A., Haas, J.L., Corriveau, E.J., Liem, A.T., Willis, K.L., Kadavy, D.R., et al. (2015) Bacterial and viral identification and differentiation by amplicon sequencing on the MinION nanopore sequencer. Gigascience 4:.

Landa, B.B. (2017) Emergence of Xylella fastidiosa in Spain: current situation.

Li, H. (2018) Minimap2: Pairwise alignment for nucleotide sequences. Bioinformatics 34: 3094–3100.

Li, W., Levy, L., Teixeira, D.C., Lopes, S., Ayres, A.J., Hartung, J.S., et al. (2013) Development and systematic validation of qPCR assays for rapid and reliable differentiation of *Xylella fastidiosa* strains causing citrus variegated chlorosis. J Microbiol Methods 92: 79–89.

Marçais, G. and Kingsford, C. (2011) A fast, lock-free approach for efficient parallel counting of occurrences of k-mers. Bioinformatics 27: 764–770.

Marcelletti, S. and Scortichini, M. (2016) Genome-wide comparison and taxonomic relatedness of multiple *Xylella fastidiosa* strains reveal the occurrence of three subspecies and a new *Xylella* species. Arch Microbiol 198: 803–812.

Marchi, G., Rizzo, D., Ranaldi, F., Ghelardini, L., Ricciolini, M., Scarpelli, I., et al. (2018) First detection of *Xylella fastidiosa* subsp. multiplex DNA in Tuscany (Italy). Phytopathol Mediterr 57: 363–364.

Modesti, V., Pucci, N., Lucchesi, S., Campus, L., and Loreti, S. (2017) Experience of the Latium region (Central Italy) as a pest-free area for monitoring of *Xylella fastidiosa*: distinctive features of molecular diagnostic methods. Eur J Plant Pathol 148: 557–566.

Nunney, L., Elfekih, S., and Stouthamer, R. (2012) The importance of multilocus sequence typing: Cautionary tales from the bacterium *Xylella fastidiosa*. Phytopathology 102: 456–462.

Olmo, D., Nieto, A., Adrover, F., Urbano, A., Beidas, O., Juan, A., et al. (2017) First Detection of *Xylella fastidiosa* Infecting Cherry (Prunus avium) and Polygala myrtifolia Plants, in Mallorca Island, Spain. Plant Dis 101: PDIS-04-17-0590.

Page, A.J. and Keane, J.A. (2018) Rapid multi-locus sequence typing direct from uncorrected long reads using Krocus. PeerJ 6: e5233.

Radhakrishnan, G. V, Cook, N.M., Bueno-Sancho, V., Lewis, C.M., Persoons, A., Mitiku, A.D., et al. (2019) MARPLE, a point-of-care, strain-level disease diagnostics and surveillance tool for complex fungal pathogens. BMC Biol 17: 1–17.

Saponari, M., Boscia, D., Nigro, F., and Martelli, G.P. (2013) Identification of dna sequences related to *Xylella fastidiosa* in oleander, almond and olive trees exhibiting leaf scorch symptoms in Apulia (Southern Italy). J Plant Pathol 95: 668.

Saponari, M., D’Attoma, G., Kubaa, R.A., Loconsole, G., Altamura, G., Zicca, S., et al. (2019) A new variant of *Xylella fastidiosa* subspecies *multiplex* detected in different host plants in the recently emerged outbreak in the region of Tuscany, Italy. Eur J Plant Pathol 1–6.

Scally, M., Schuenzel, E.L., Stouthamer, R., and Nunney, L. (2005) Multilocus sequence type system for the plant pathogen *Xylella fastidiosa* and relative contributions of recombination and point mutation to clonal diversity. Appl Environ Microbiol 71: 8491–8499.

Schaad, N.W., Opgenorth, D., and Gaush, P. (2002) Real-time polymerase chain reaction for one-hour on-site diagnosis of Pierce’s disease of grape in early season asymptomatic vines. Phytopathology 92: 721–728.

Sherald, J.L. and Kostka, S.J. (1992) Bacterial Leaf Scorch of Landscape Trees Caused By Xylella fastidiosa 1. 18: 57–63.

Vaser, R., Sović, I., Nagarajan, N., and Šikić, M. (2017) Fast and accurate de novo genome assembly from long uncorrected reads. Genome Res 27: 737–746.

Votintseva, A.A., Bradley, P., Pankhurst, L., Del Ojo Elias, C., Loose, M., Nilgiriwala, K., et al. (2017) Same-day diagnostic and surveillance data for tuberculosis via whole-genome sequencing of direct respiratory samples. J Clin Microbiol 55: 1285–1298.

Vyahhi, N., Prjibelski, A.D., Nurk, S., Pyshkin, A. V., Dvorkin, M., Alekseyev, M. a., et al. (2012) SPAdes: A New Genome Assembly Algorithm and Its Applications to Single-Cell Sequencing. J Comput Biol 19: 455–477.

Wells, J.M., Raju, B.C., Hung, H.-Y., Weisburg, W.G., Mandelco-Paul, L., and Brenner, D.J. (1987) *Xylella fastidiosa* gen. nov., sp. nov: Gram-Negative, Xylem-Limited, Fastidious Plant Bacteria Related to Xanthomonas spp. Int J Syst Bacteriol 37: 136–143.

Wick, R.R., Judd, L.M., and Holt, K.E. (2018) Deepbinner: Demultiplexing barcoded Oxford Nanopore reads with deep convolutional neural networks. PLoS Comput Biol 14:.

Wick, R.R., Judd, L.M., and Holt, K.E. (2019) Performance of neural network basecalling tools for Oxford Nanopore sequencing. Genome Biol 20: 129.

Yuan, X., Morano, L., Bromley, R., Spring-Pearson, S., Stouthamer, R., and Nunney, L. (2010) Multilocus Sequence Typing of *Xylella fastidiosa* Causing Pierce’s Disease and Oleander Leaf Scorch in the United States. Phytopathology 100: 601–611.

